# Promoting Longevity in Aged Liver through NLRP3 Inflammasome Inhibition Using Tauroursodeoxycholic Acid (TUDCA) and SCD Probiotics

**DOI:** 10.1101/2024.02.27.582399

**Authors:** Burcu Baba, Taha Ceylani, Eda Acikgoz, Rafig Gurbanov, Seda Keskin, Gizem Samgane, Huseyin Tombuloglu, Hikmet Taner Teker

## Abstract

This study investigates the combined impact of SCD Probiotics and tauroursodeoxycholic acid (TUDCA) on the biomolecular makeup, histological changes and levels of inflammasome in the liver tissue of 24-month-old male Sprague-Dawley rats. By administering TUDCA (300 mg/kg, intravenously) and SCD Probiotics (3 mL (1 x 108 CFU), orally) daily for a week, the researchers employed ATR-FTIR spectroscopy along with machine learning approaches such as Linear Discriminant Analysis (LDA) and Support Vector Machine (SVM) to analyze the biomolecular profiles. In addition, the study measured the expression levels of inflammasome markers NLRP3, ASC, Caspase-1, IL18, and IL1β using RT-qPCR and examined liver sections for histopathological changes and NLRP3 inflammasome activation. The results revealed significant differences in the levels of lipids, proteins, and nucleic acids, with TUDCA having a noteworthy impact on enhancing lipid bands and reducing cholesterol ester bands, while SCD Probiotics showed the opposite effects. Furthermore, TUDCA was found to decrease the acyl chain length of fatty acids and improve protein conformation, whereas SCD Probiotics increased both the acyl chain length and protein phosphorylation ratio, suggesting a decrease in lipid and protein dynamics from both treatments. The histological assessments showed significant reductions in cellular degeneration, lymphatic infiltration, hepatic fibrosis, and the immunoreactivity of NLRP3 and ASC in the treated groups. SCD Probiotics exhibited a marked reduction in inflammasome-related gene expressions, and the lowest gene expression levels were observed in the group receiving both treatments. Despite an increase in serum AST and LDH levels across all groups, only the SCD Probiotics group showed an increase in albumin levels. The findings suggest that SCD Probiotics, TUDCA, and their combined administration may provide a promising avenue for therapeutic interventions in age-associated liver conditions and may mitigate age-related liver fibrosis while enhancing liver functionality.

## INTRODUCTION

Probiotics are living microorganisms that, when consumed in adequate amounts, provide health benefits. Among their many benefits, probiotics have been shown to support healthy aging, rejuvenation, and liver health (1). Changes occur in the microbiome with aging, including a decline in the abundance of beneficial bacteria (2). This shift is associated with the onset of age-related diseases, including metabolic disorders, cardiovascular disease, and inflammation. Probiotics have been found to improve gut microbiota, promote immune function, and decrease inflammation, which can help to reduce the risk of these diseases (3). The liver is a vital organ that plays a critical role in metabolism, detoxification, and energy storage. However, liver function declines with age, which can lead to the development of liver diseases such as non-alcoholic fatty liver disease (NAFLD). Studies have shown that probiotics can aid in the removal of toxins from the liver and prevent various liver health issues, including NAFLD (4). Furthermore, probiotics’ anti-inflammatory properties can prevent liver inflammation (5). In a randomized controlled trial, probiotic supplementation was found to improve liver function and reduce the risk of liver disease in elderly participants (6). The positive effects of probiotics on gut microbiota, immune function, liver health, and skin rejuvenation make them an important tool for healthy aging. Therefore, incorporating probiotics into the diet may be an essential component of maintaining good health in old age (7).

Bile acids are molecules that are produced in the liver from cholesterol, possessing both hydrophilic and hydrophobic properties, and serving a critical role in lipid digestion. The gut microbiota regulates the synthesis and metabolism of bile acids (8). Tauroursodeoxycholic acid (TUDCA) is a type of conjugated bile acid derivative formed by the combination of ursodeoxycholic acid (UDCA) and taurine. The gut microbiota generates UDCA, which then returns to the liver via enterohepatic circulation, where it conjugates with taurine to create TUDCA (9). TUDCA has been shown to improve the gut microbiota (10) and reduce inflammation and barrier disruption in the intestines in animal studies (11,12). TUDCA has a wide range of health benefits, particularly for liver and aging health. It is believed to work by protecting liver cells from toxic bile acids, reducing inflammation, and regulating energy metabolism, which can enhance liver health (13). Additionally, TUDCA improves insulin sensitivity, which is essential for maintaining metabolic health and preventing diabetes (14). Its potential to safeguard against age-related diseases makes it a promising candidate for future anti-aging therapies.

Aging is a critical process characterized by the loss of hepatic function due to the accumulation of reactive oxygen species (15). Age-related inflammation disrupts interactions within the inflammasome signaling pathway, which serves a vital function within the cell. The NLRP3 inflammasome can be overactivated and produce an aggravated inflammatory process after chronic damage, such as metabolic disorders occurring during liver fibrosis and steatosis, or during a progressive deterioration process associated with aging (16). The relationship between the NLRP3 inflammasome complex and age-related liver damage is mediated by the altered expression of NLRP3 inflammasome components and the eventual activation of an inflammatory process in the liver. Therefore, the progression of severe activation of inflammasomes in this molecular framework can be defined as a crucial trigger in aging (17).

This study aims to investigating the synergistic effects of SCD Probiotics and TUDCA on histopathology and biomolecular profiles in aged rat liver tissue. By examining the biomolecular structures, histological alterations, gene exspression and employing advanced analytical techniques, we hope to shed light on the potential benefits of these treatments in mitigating age-related liver damage. The insights gained from this study could pave the way for further investigations into the therapeutic potential of SCD Probiotics and TUDCA in managing age-related liver disorders and improving the quality of life particularly for the elderly population.

## MATERIALS AND METHODS

This section is presented as a supplementary file.

## RESULTS

### TUDCA and SDC Probiotics caused significant changes in liver biomolecules

Significant qualitative modifications were detected by LDA analysis with 100% accuracy for full biomolecules of the liver (**Fig. 1, Tables S1-S2**). Based on the discrimination plot, the data for the control (CLI), SDC Probiotics (PLI), TUDCA (TLI), and the applications (PTLI) in which the TUDCA and SCD Probiotics were applied together are clustered in separate coordinates (**Fig. 1**). The same qualitative changes were also seen for the lipids, proteins, and nucleic acids with 100% accuracy (**Fig. 2 and Figs. S1-S2, Tables S3-S8**). SVM classification capacity was also quite evident for the whole biomolecular content of the liver, revealing high training (95%) and validation accuracies (92.5%) **(Fig. S3).**

**Fig. 1.**
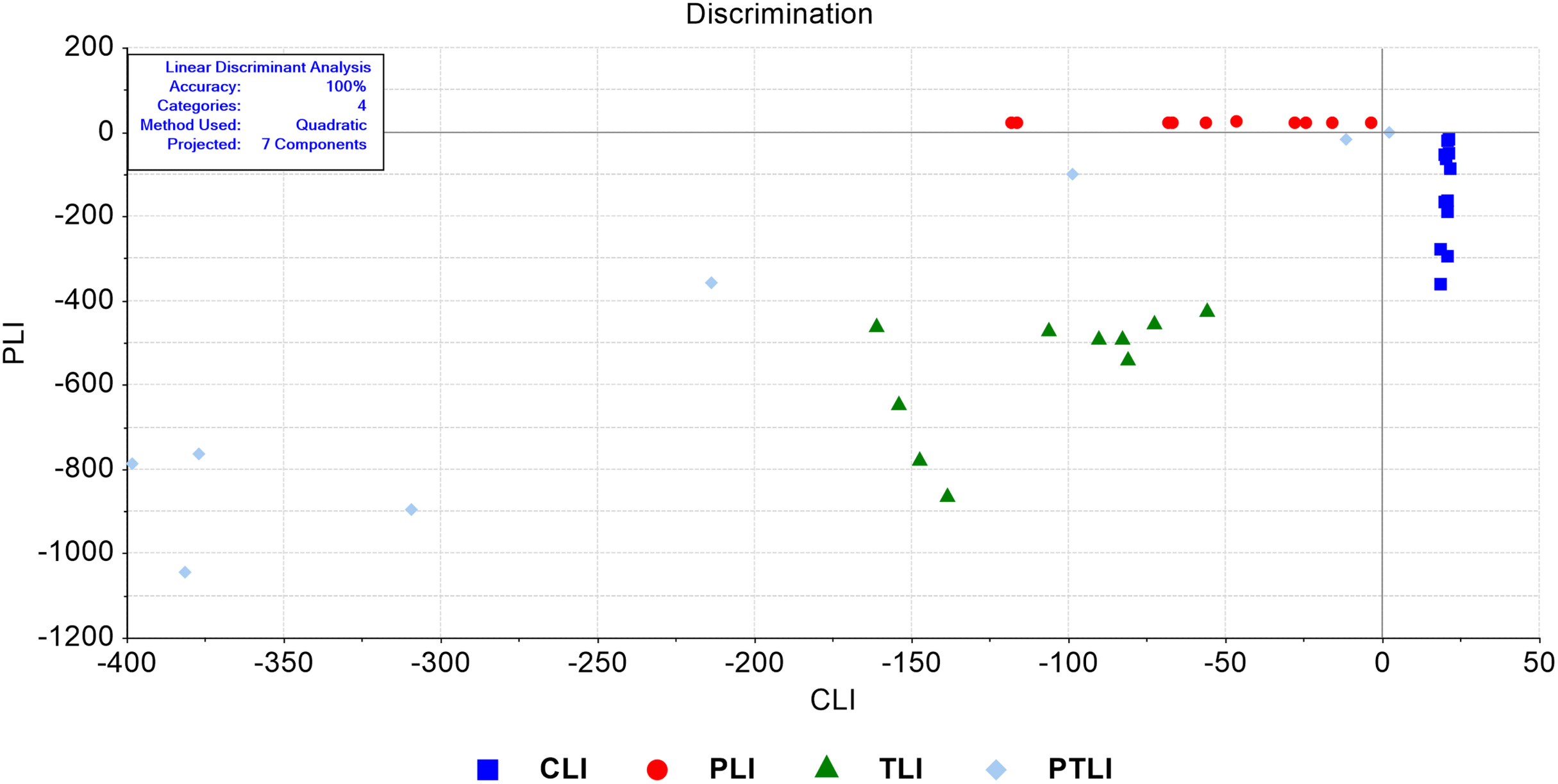
LDA discrimination plot for liver samples in the full (4000-650 cm^-1^) spectral region. CLI (control), TLI (TUDCA), PLI (SDC Probiotics), and the PTLI applications (in which the TUDCA and SCD Probiotics were applied together)

**Fig. 2.**
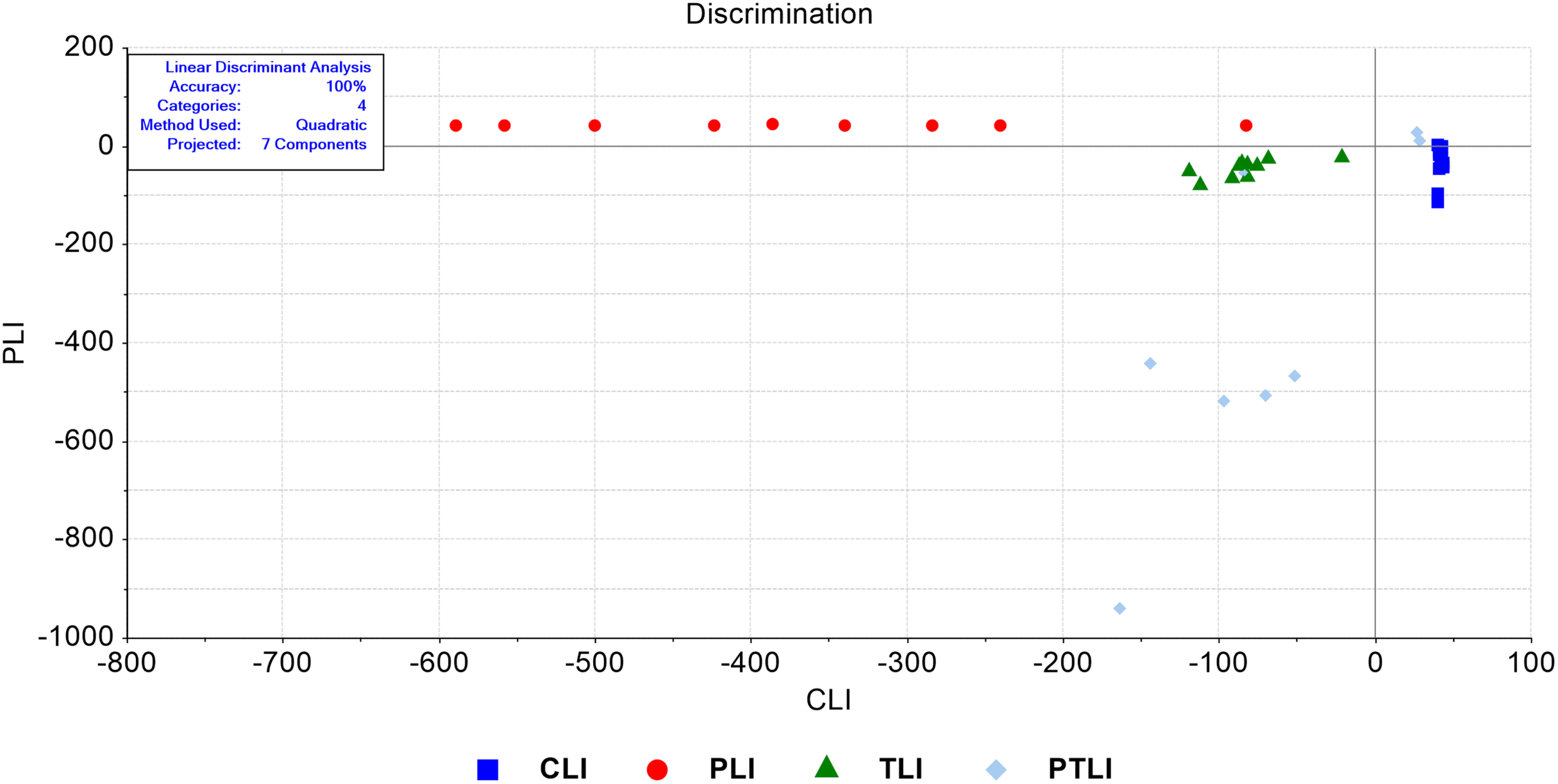
LDA discrimination plot for liver samples in the lipid (3000-2700 cm^-1^) spectral region. CLI (control), TLI (TUDCA), PLI (SDC Probiotics), and the PTLI applications (in which the TUDCA and SCD Probiotics were applied together)

The quantitative analyses of spectral bands at 2955 cm^-1^ (CH_3_ antisymmetric stretching: lipids and proteins), 2922 cm^-1^ (CH_2_ antisymmetric stretching: lipids), 1740 cm^-1^ (C=O stretching: cholesterol ester), 1653 cm^-1^ (Amide I: α*-*helical structure of proteins), 1545 cm^-1^ (Amide II: β*-*sheet structure of proteins), 1239 cm^-1^ (PO_2_ antisymmetric stretching: nucleic acids), and 1080 cm^-1^ (PO_2_ symmetric stretching: nucleic acids) positions were performed for detailed analysis of biomolecules (**Fig. 3a-f**). The CH3 antisymmetric stretching, a commonly used lipid marker, increased in the TUDCA group but decreased in the SCD Probiotics group. The CH2 antisymmetric, another lipid marker, decreased in all groups, with the greatest reduction observed in the TUDCA group. The C=O stretching, a marker of cholesterol ester, increased in the SCD Probiotics group but decreased in the TUDCA group (**Fig. 3a-c**). The Amid I band, a widely used protein marker, did not show a significant difference between groups. However, in the group where both applications were used, there was a significant increase in the Amid II band, which is also considered a protein marker. (**Fig. 3d**). TUDCA caused a significant decrease in the PO2 antisymmetric and PO2 symmetric bands, which are considered nucleic acid markers, while SDC Probiotics only resulted in a significant decrease in the PO2 antisymmetric band (**Fig. 3e-f**).

**Fig. 3.**
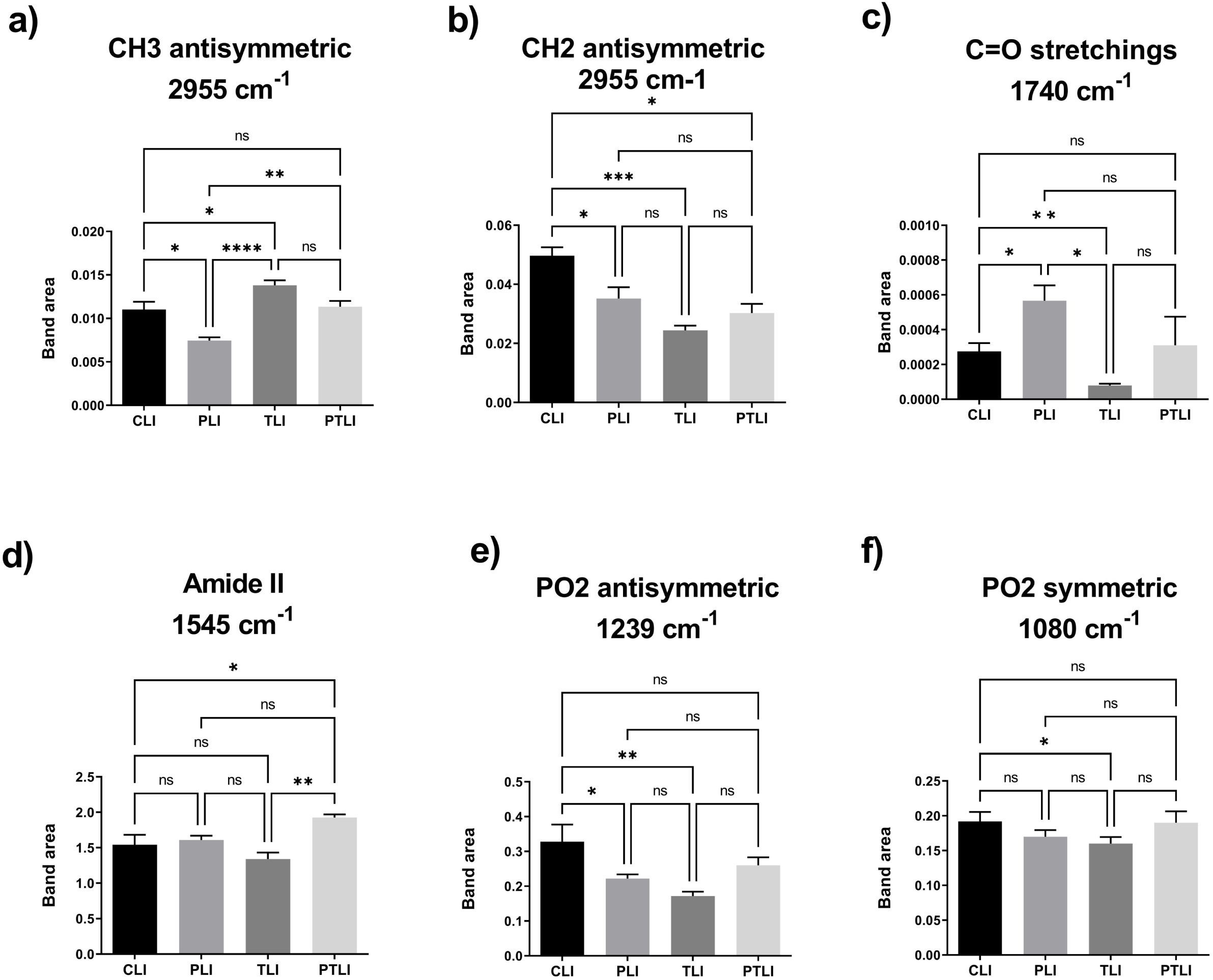
The quantitative changes in liver-associated spectrochemical parameters. The band areas for **a)** CH_3_ antisymmetric (2955 cm^-1^), **b)** CH_2_ antisymmetric (2922 cm^-1^), **c)** lipid carbonyl (C=O stretchings, 1740 cm^-1^), **d)** Amide II (1545 cm^-1^), **e)** PO2 antisymmetric (1239 cm-1), and **f)** PO_2_ symmetric (1080 cm^-1^). CLI (control), TLI (TUDCA), PLI (SDC Probiotics), and the PTLI applications (in which the TUDCA and SCD Probiotics were applied together)

The band area ratios for the acyl chain of fatty acids (A_2922_/A_2955_), Protein phosphorylation (A_1239_/A_2955_), Protein phosphorylation (A_1080_/A_1545_), Protein conformation (A_1653_/A_1545_) and Protein carbonylation (A_1740_/A_1545_) of proteins were changed significantly (**Fig. 4a-d**). The SCD Probiotics group showed an increase in the acyl chain of fatty acids, whereas the TUDCA group showed a decrease (**Fig. 4a**). Similarly, the phosphorylation ratio (A_1239_/A_2955_) increased in the SCD Probiotics group but decreased in the TUDCA group. However, the phosphorylation ratio (A_1080_/A_1545_) only increased in the TUDCA group (**Fig. 4b-c**). Protein conformation (A_1653_/A_1545_) analysis showed a significant increase only in the TUDCA group (**Fig. 4d**). Protein carbonylation significantly decreased in all treated group (**Fig. 4e**).

**Fig. 4.**
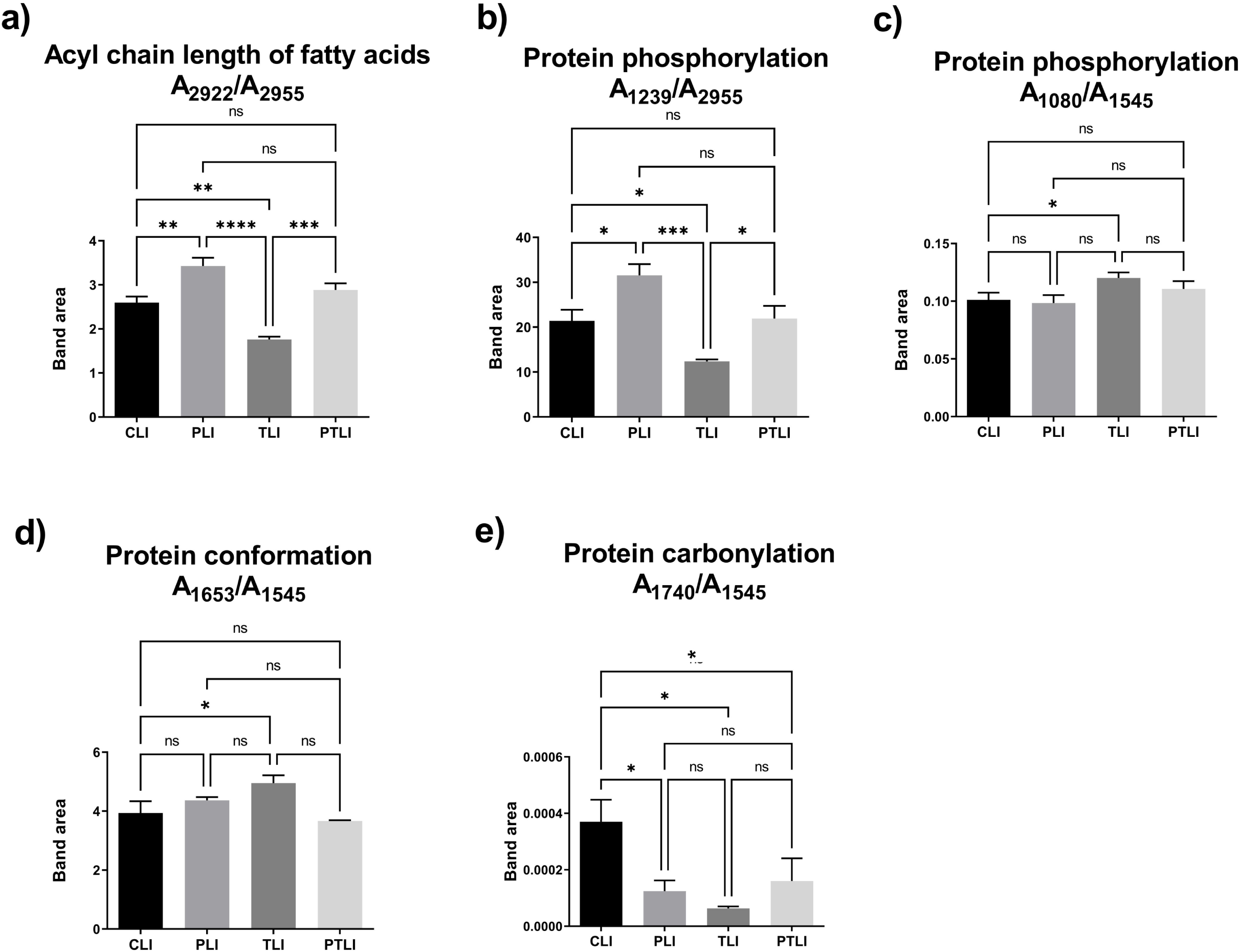
The quantitative changes in liver-associated spectrochemical parameters. The band area ratios for **a)** acyl chain length of fatty acids (A_2922_/A_2955_), **b)** protein phosphorylation (A_1239_/A_2955_), **c)** protein phosphorylation (A_1239_/A_2955_), **d)** protein conformation (A_1653_/A_1545_), and **e)** protein carbonylation (A_1740_/A_1545_). CLI (control), TLI (TUDCA), PLI (SDC Probiotics), and the PTLI applications (in which the TUDCA and SCD Probiotics were applied together)

Moreover, bandwidths of spectral parameters 2955 cm^-1^ (CH3 antisymmetric stretching: lipids and proteins), 2922 cm^-1^ (CH2 antisymmetric stretching: lipids), 1653 cm^-1^ (Amide I: α-helical structure of proteins), and membrane dynamics (A_2922_/A_2955_) changed significantly (**Fig. S4a-d**). The CH3 antisymmetric band increased only in the TUDCA group, while the CH2 antisymmetric, Amide I, and membrane dynamics decreased in both the SCD Probiotics and TUDCA groups.

### Effects of SCD Probiotics, TUDCA and combined theatment on age-related liver changes

H&E staining was performed to evaluate the histological changes in aged liver due to SCD probiotics, TUDCA, and combined treatments. As shown in **Fig. 5a**, irregular hepatocyte arrangement, enlargement of sinusoidal areas, relative increase in bile duct density and Kupffer cells, and intense lymphatic and neutrophil infiltration were observed in CLI group. On the other hand, PLI, TLI and combined treatment groups showed a significant decrease in lymphocytic and neutrophil infiltration intensity and improvements in cellular degenerations developing in the aging process compared to the CLI group. From the findings obtained, it was observed that the combined treatment of SCD Probiotics and TUDCA may educe age-related inflammation and bile duct proliferation in the aged liver. However, when the effects of SCD Probiotics, TUDCA and combined treatment on microvesicular steatosis, a type of hepatic fat accumulation in aged livers were examined. It was observed that microvesicular steatosis was significantly increased in the CLI group, whereas a significant decrease in the density of lipid droplets was observed in the SCD Probiotics, TUDCA and combine treated groups compared to the CLI group (**Fig. 5b**). There was no difference in the improvement rate (steatosis) between PLI only and TLI only groups. All findings indicate the potential of SCD Probiotics and TUDCA treatments to alleviate age-related liver damage, inflammation and lipid metabolism disorders.

**Fig 5.**
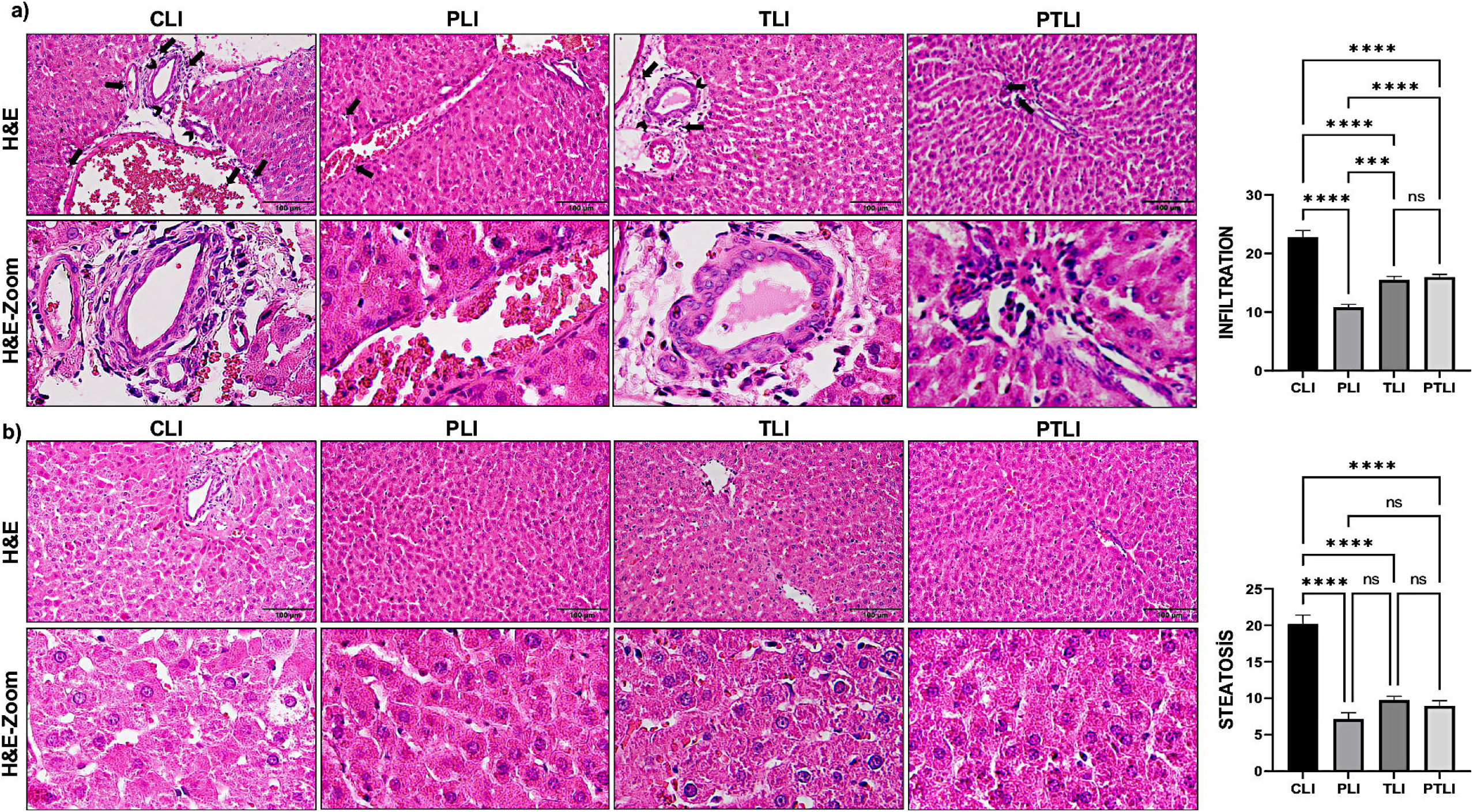
Effects on liver histological architecture **a.** Representative images of hematoxylin and eosin (H&E) staining and quantification of infiltration **b.** Representative images of H&E staining and quantification of micro vesicular steatosis. Black arrows show inflammation. Black arrow’s heads indicate bile ducts. The areas in the relevant microphotographs of all groups are enlarged below. Values are expressed as mean ±SEM. p≤ 0.001 ***, and p≤ 0.0001 ****, ns: not significant. Scale bar:100 µm. CLI (control), TLI (TUDCA), PLI (SDC Probiotics), and the PTLI applications (in which the TUDCA and SCD Probiotics were applied together).

### SCD Probiotics, TUDCA and combined treatments alleviates liver fibrosis in the aged liver

Collagen accumulation density changes in MT stained liver sections were analyzed by ImageJ to investigate whether SCD Probiotics, TUDCA and combined treatments alleviate age-related liver fibrosis. In the results, the intensities of blue stained areas representing collagen accumulation in the liver tissue of the PLI, TLI, and PTLI groups were significantly decreased compared to the CLI group **(Fig. 6).** These results together suggest that SCD probiotics, TUDCA, and combined treatment could reduce age-related liver fibrosis.

**Fig 6.**
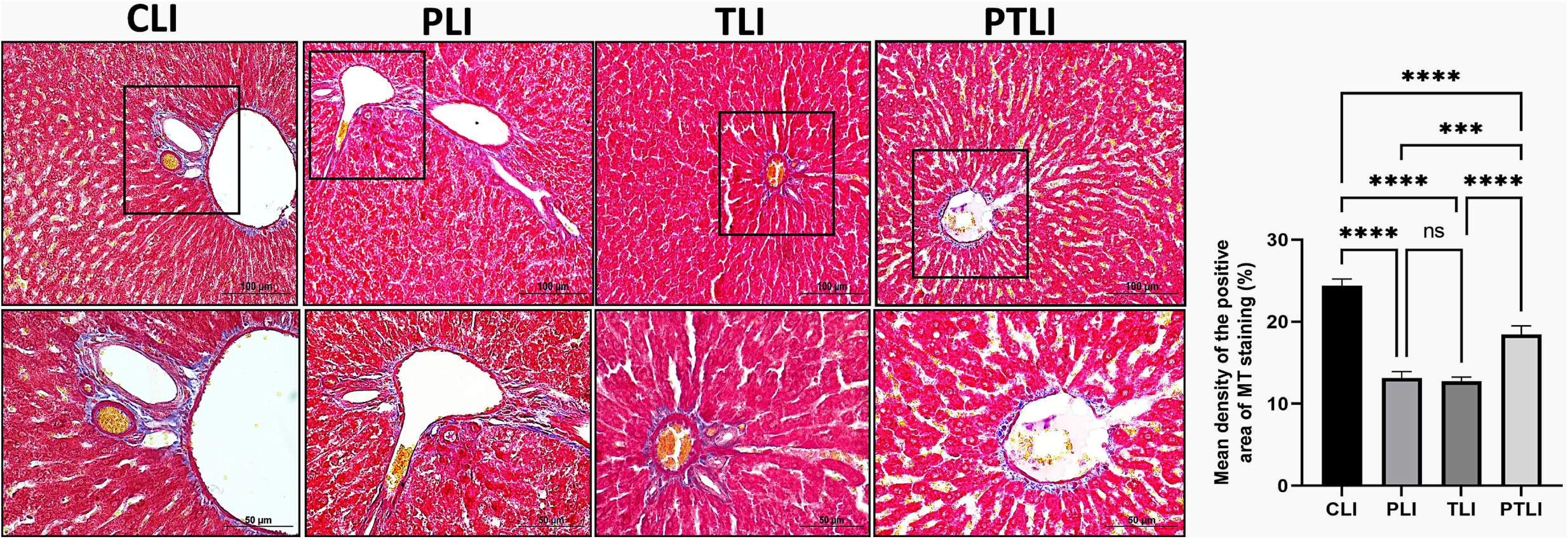
Effects on liver fibrosis. Representative images of Masson trichrome (MT) staining showing collagen deposition in the aged contol rat and all treatment groups liver tissue with quantification of collagen density area fraction (%) in all groups. The areas enclosed in squares in the relevant microphotographs of all groups are enlarged below. Values are expressed as mean ±SEM. p≤ 0.001 ***, and p≤ 0.0001 ****, ns: not significant. Scale bar:50 and 100 µm. CLI (control), TLI (TUDCA), PLI (SDC Probiotics), and the PTLI applications (in which the TUDCA and SCD Probiotics were applied together)

### SCD Probiotics and TUDCA exhibited a marked reduction in inflammasome-related gene expressions

The results of the study involving the use of SCD Probiotics, Tauroursodeoxycholic acid (TUDCA), and their combination in modulating the expression of NLRP3, ASC, Caspase-1, IL18, and IL1β genes related to inflammasomes are presented. The application of SCD Probiotics alone demonstrated the most significant reduction in the expression of the genes, as shown in **Figure 7**. Interestingly, the combination of SCD Probiotics and TUDCA resulted in a less pronounced effect on gene expression levels compared to when each treatment was applied independently. These observations were further supported by Immunohistochemistry (IHC) staining, which reinforced the initial findings and provided a deeper understanding of the molecular mechanisms underlying the anti-inflammatory effects of the treatments.

**Fig. 7.**
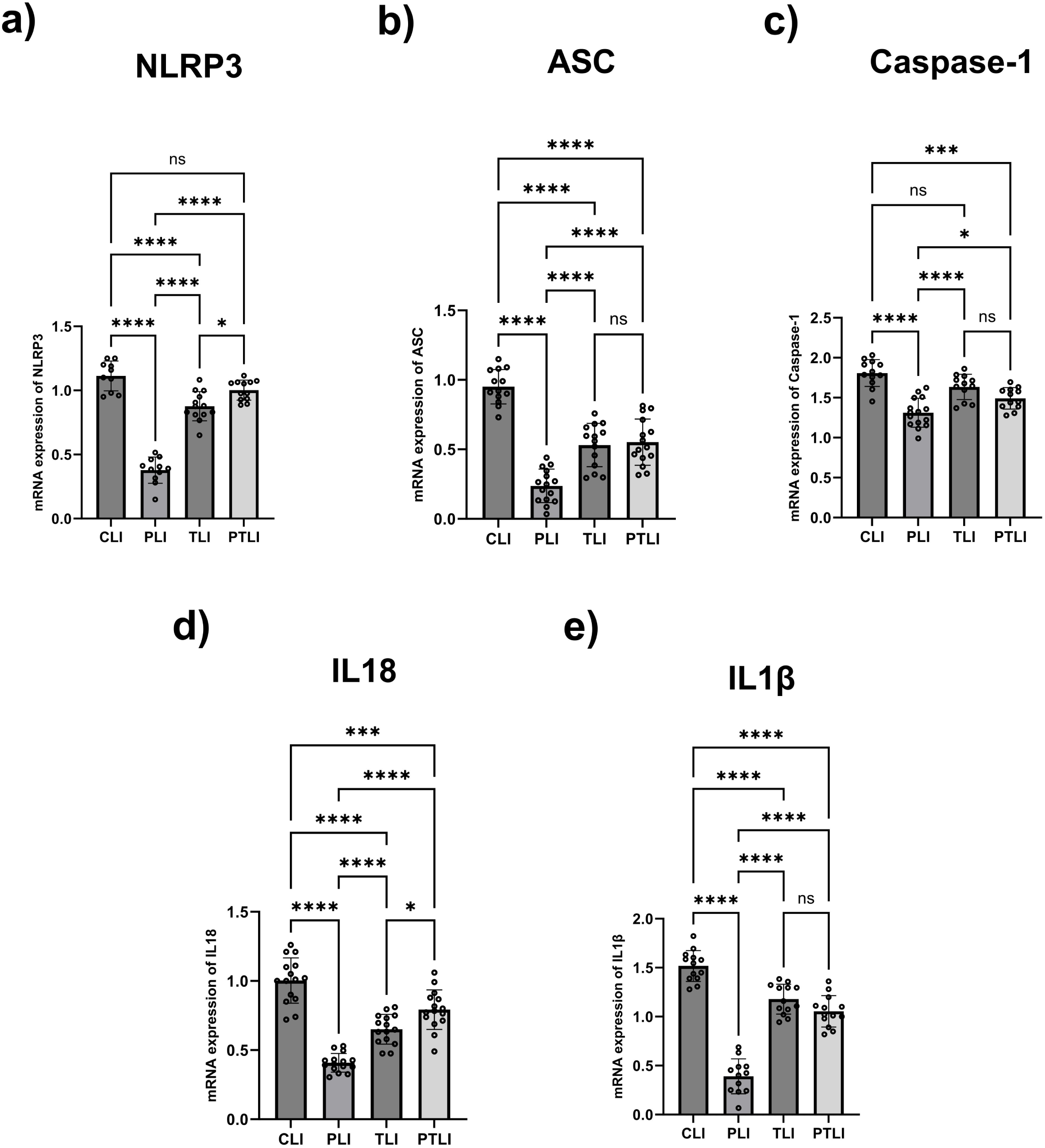
The relative mRNA expression levels in inflammasome-related gene expressions. (A) mRNA expression levels of NLRP3, (B) mRNA expression levels of ASC, (C) mRNA expression levels of Caspase-1, (D) mRNA expression levels of IL18, and (E) mRNA expression levels of IL1β. All data are shown as Mean ± SEM (n = 5 and 3 replicates); p values are derived using Tukey’s Multiple Comparisons Test followed by One-way ANOVA. Statistically significant differences are indicated in the following way: ns p > 0.05 (not significant); * p < 0.05 (significant); *** p < 0.001 and **** p < 0.0001 (extremely significant). CLI (control), TLI (TUDCA), PLI (SDC Probiotics), and the PTLI applications (in which the TUDCA and SCD Probiotics were applied together)

### SCD Probiotics and TUDCA administered to aged rats reduce inflammation by suppressing NLRP3 inflammasome activation in liver tissue

Hyperactivation of the NLRP3 inflammasome contributes to inflammatory responses and cellular damage in aging (18). Our histological results revealed severe inflammation in the aged liver. Therefore, expression levels of NLRP3 and ASC associated with the NLRP3 inflammasome complex were analyzed using IHC-stained liver sections. As shown in **Fig. 7**, NLRP3 and ASC expression were found to be high, particularly in areas of intense inflammation. Compared with the CLI group, a decrease in the expression levels of NLRP3 and ASC was seen in the PLI, TLI, and PTLI groups (p<0.05). Furthermore, there was no statistical difference in NLRP3 and ASC expression levels between the TLI and combination groups (p>0.05). Together, the results obtained showed that NLRP3 inflammasome activation triggered by inflammation in the aged liver was significantly suppressed by SCD Probiotics, TUDCA and combined treatments.

### Biochemical analysis of liver enzymes in blood serum

The results showed that serum AST levels significantly increased in the treatment groups compared to the control group **(Fig. 8a)**, while ALT and ALP levels showed no significant differences. AST, not the only liver-specific enzyme, is also found in muscle tissue, which decreases in elderly individuals due to cellular degeneration. Consequently, AST levels may be low in aged rats. LDH, an essential enzyme found primarily in muscle, liver, and kidney cells, showed increased serum levels in the treatment groups compared to the control group **(Fig. 8b)**. Metabolic adaptations associated with aging are not well understood, but changes in albumin levels have been linked to acute and chronic liver diseases. A decrease in serum albumin concentration with age has been previously reported, but this study found that treated groups had significantly increased serum albumin levels **(Fig. 8c)**.

**Fig 8.**
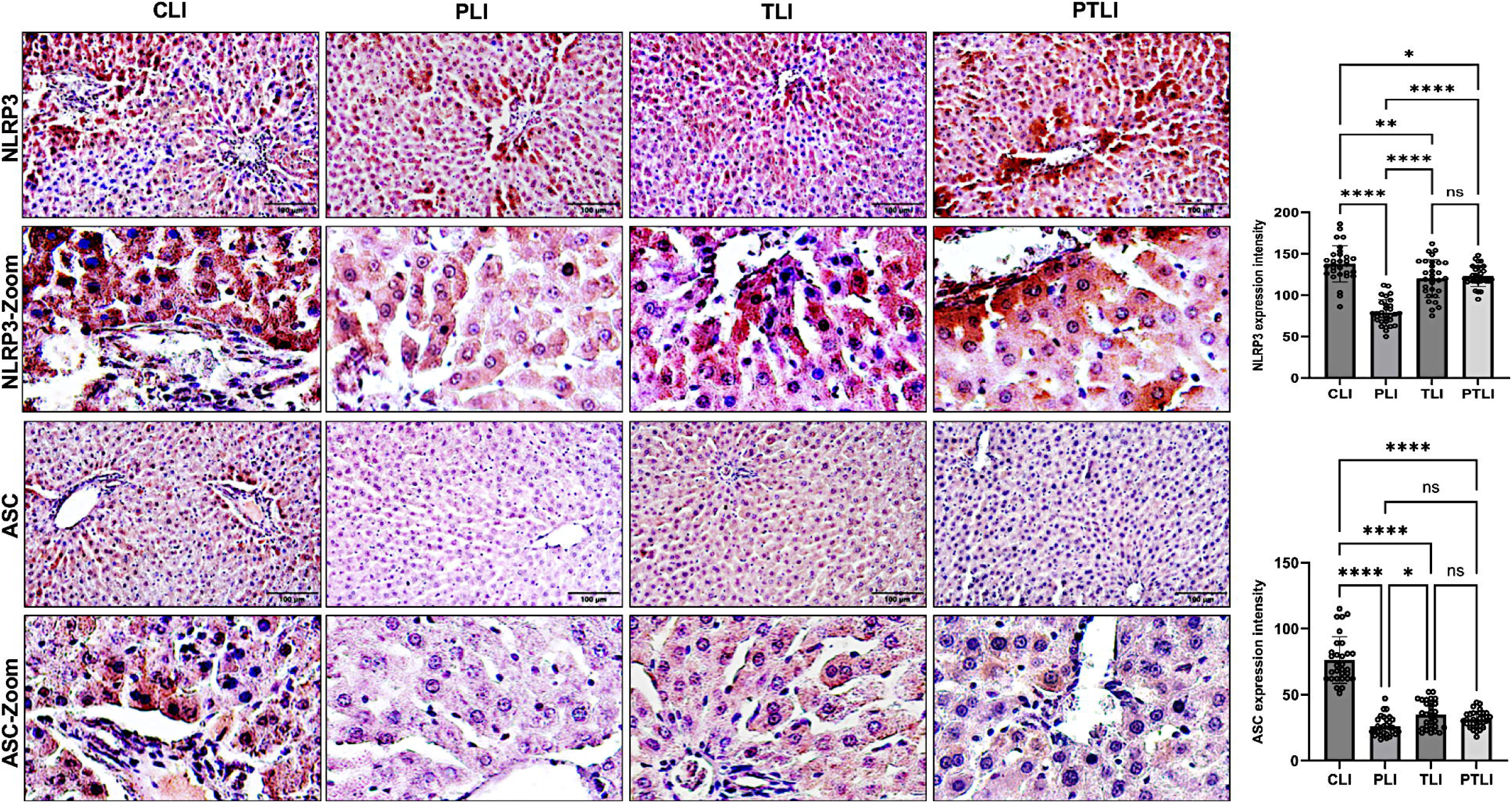
Representative photomicrographs showing NLRP3, and ASC immunostaining and respective quantifications of expression intensity in the liver. The zooms of the selected areas are shown below the respective microphotographs. Expression intensities (NLRP3 and ASC) are given in the right panel. Values are expressed as mean ± SEM. p<0.05 *, p≤ 0.01 **, and p≤ 0.0001 ****, ns: not significant. Scale bar: 100 µm. CLI (control), TLI (TUDCA), PLI (SDC Probiotics), and the PTLI applications (in which the TUDCA and SCD Probiotics were applied together)

**Fig 9.**
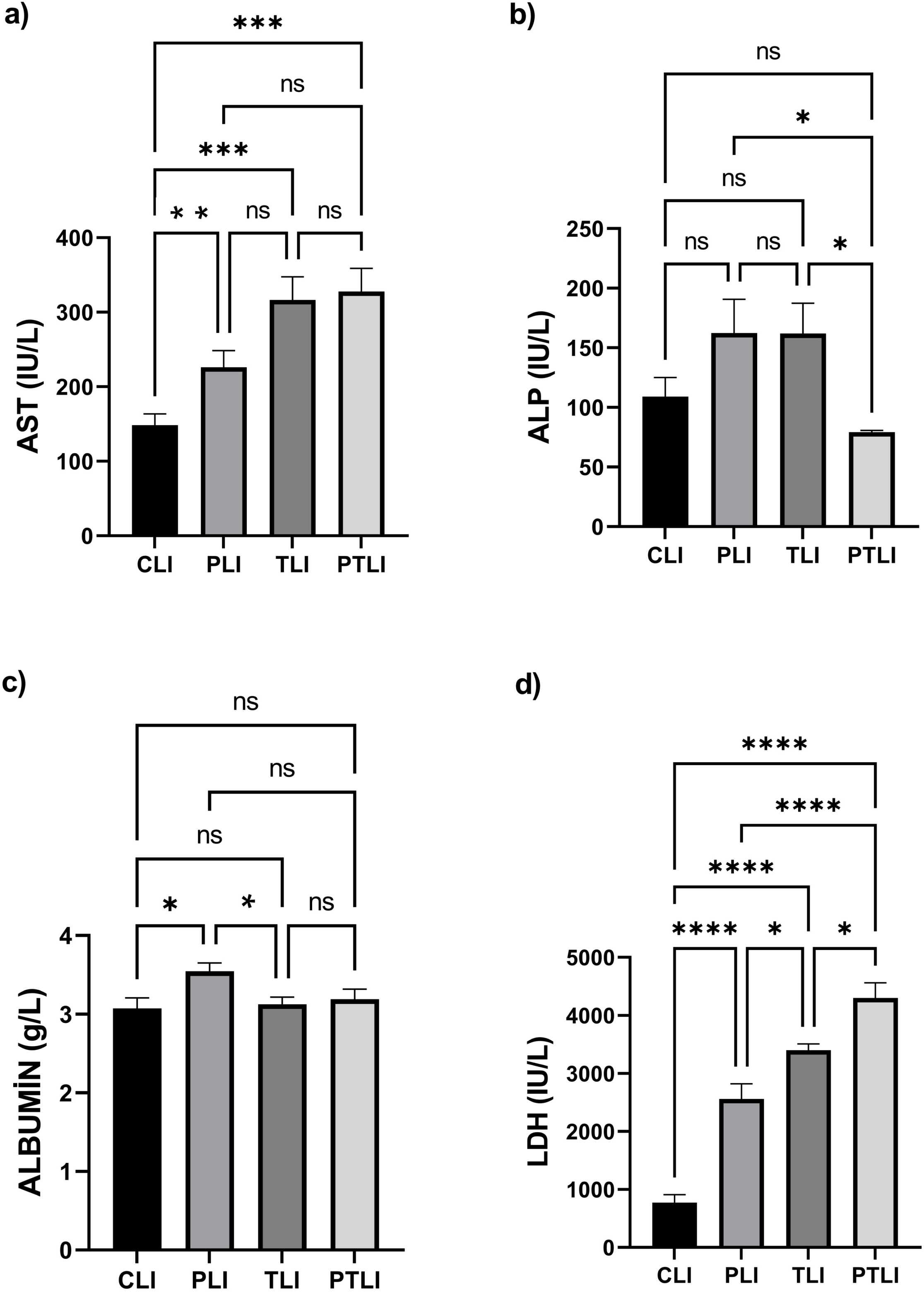
Effects of all treatment groups (PLI, TLI and PTLI groups) on the serum levels of **a)** AST, **b)** ALP, **c)** Albumin, and **d)** LDH for one week treatment. Values were provided as meanL±LSD, nL=L7/group. *PL<L0.05 and **PL<L0.01 vs. control. CLI (control), TLI (TUDCA), PLI (SDC Probiotics), and the PTLI applications (in which the TUDCA and SCD Probiotics were applied together)

## DISCUSSION

In the process of aging, the liver undergoes significant transformations, including a decline in blood flow and regenerative capacity, accumulation of fat, decreased detoxification abilities, and gene expression alterations (19,20). These shifts can boost the risk of liver disease and other health issues in seniors. The aging-related imbalance in gut microbiota, known as dysbiosis, can also lead to a decline in bacterial diversity, weakened gut barrier, and various metabolic disorders (21,22). This study explores how a combination of SCD Probiotics and TUDCA impacts the liver’s molecular and tissue structure in aged rats. A machine learning approach and spectroscopy were used to examine molecular profiles, demonstrating distinct effects of TUDCA and SCD Probiotics on lipid, protein, and nucleic molecules. To better understand these differences, infrared (IR) spectroscopy was used (23). Yet, when TUDCA and SCD Probiotics were used together, some effects were less pronounced or contradictory, complicating conclusions about their collective or separate benefits. For example, while TUDCA increased a lipid marker (CH3 antisymmetric band), SCD Probiotics reduced it, suggesting possible lipid peroxidation in the liver, which could be negative (24). However, free radicals, sometimes increased by such changes, play important roles in physiological signaling and probiotics can reduce oxidative damage in liver disease patients (25,26).

Aging can impact liver’s cholesterol metabolism, leading to increased accumulation. TUDCA, in this study, enhanced bile acid synthesis in the liver, and reduced cholesterol build-up in serum and liver of mice, as marked by a decrease in the C=O stretching band, a cholesterol ester marker (10). Contrarily, SCD Probiotics increased this band, contrary to previous research indicating that these probiotics, containing strains like *B. longum*, *L. acidophilus*, and *L. plantarum*, usually reduce cholesterol levels in serum and liver (27). Further, TUDCA caused a decrease, while SCD Probiotics led to an increase in the acyl chain of fatty acids band, a component of lipids involved in membrane structure and function, energy storage, and cellular signaling (28). While an increase can be seen as beneficial, saturated lipid acyl chains could create a less fluid bilayer, so a decrease might be advantageous. This decrease with TUDCA aligns with recent studies suggesting its role in modulating lipid metabolism, possibly by controlling the synthesis and breakdown of fatty acids and other lipids (29). However, in the group receiving both treatments, no significant difference was seen in these parameters, suggesting further analyses are necessary to draw clear conclusions.

The endoplasmic reticulum (ER) is a complex organelle that plays a crucial role in various cellular processes, including protein synthesis, folding, and modification, lipid metabolism, and calcium homeostasis. Research studies suggest that TUDCA can attenuate ER stress, prevent dysfunction in the unfolded protein response, and stabilize mitochondria (30). In spectrochemical analysis of proteins, it has been observed that there is a significant difference in protein conformation only in the group treated with TUDCA, indicating its effect. Furthermore, TUDCA appears to be more effective in terms of protein phosphorylation. However, the interpretation of the decrease or increase in protein phosphorylation levels requires further analysis. Protein phosphorylation is a process where a phosphate group is added to a protein molecule by a protein kinase enzyme. This modification can regulate the activity of the protein, often by activating or deactivating it. Protein phosphorylation plays a crucial role in a variety of cellular processes, including signal transduction, gene expression, and cell cycle control. It is also involved in various diseases, such as cancer, diabetes, and neurodegenerative disorders (31). Another important parameter in which TUDCA was more effective was membrane dynamics. Interestingly, both SCD Probiotics and TUDCA caused a significant reduction in this band, but there was no significant difference when applied together as markers evaluated for nucleic acid.

Previous research has shown that age-related hyperinsulinemia can increase fat deposition, adipocyte size, and lymphatic infiltration, leading to fat accumulation in the liver and systemic insulin resistance (32,33). Our study suggests that the increase in microvesicular steatosis and lymphatic infiltration in control aged mice may be associated with a decrease in age-related hyperinsulinemia (34,35). In the elderly control group, we observed significantly increased microvesicular steatosis and lymphatic infiltration compared to all treatment groups. We propose that the decrease in microvesicular steatosis and lymphatic infiltration rates in treatment groups given SCD Probiotics and TUDCA could be related to age-related hyperinsulinemia, as we did not observe changes in food intake during the one-week treatment period. Furthermore, we believe that the reduction in lymphatic infiltration rates in treatment groups improves inflammation in hepatocytes due to aging, as a result of the decrease in fat deposits in the aged liver. TUDCA treatment has been previously found to effectively reduce body weight, fat content, and cellular inflammation in aged mice, as well as in diet-induced obese mice (36). Additionally, TUDCA has been reported to decrease lipid accumulation in the liver of obese mice by improving insulin sensitivity (37). Together, our study’s findings suggest that TUDCA and SCD Probiotics may be effective in preventing or reducing the severity of tissue damage in aging-related liver diseases in aged rat liver.

Studies have shown that insulin resistance is associated with triglyceride accumulation and fatty acid intermediate metabolites. In our study, the shortened acyl chain fatty acids in the elderly control group indicate the presence of insulin resistance. The increase in acyl chain fatty acid length in treatment groups given SCD Probiotics and TUDCA suggests a decrease in age-related hyperinsulinemia. This finding is closely related to the reduction of microvesicular steatosis in the liver tissue of the treatment groups and is consistent with the results of Zangerolamo et al. (2022b). Therefore, the observed reductions in hepatic triglyceride and cholesterol levels in aged rats treated with TUDCA may have contributed to a decrease in lipid droplet density in the aged liver.

Fibrosis is a sign of aging in various organs, including the liver, heart, and kidney. In liver fibrosis that develops with aging, there is an increase in the accumulation of collagen, one of the extracellular matrix elements (ECM) (39). In our study, we found a significant increase in collagen density in hepatic tissue in the MT staining results of liver tissues of aged rats. In contrast, there was a significant decrease in collagen densities in SCD Probiotics, TUDCA, and combined treatment groups. Based on the MT staining results, we concluded that SCD Probiotics and TUDCA applications can prevent liver fibrosis due to aging. In one study, when liver enzyme results were evaluated together with histopathological examinations, it was reported that TUDCA treatment significantly improved serum ALT, AST, ALP, and albumin concentrations. Among patients who underwent percutaneous liver biopsy, a histological relief of tissue fibrosis in the liver was observed in one patient in the TUDCA group (40). In another study, it is believed that exogenous administration of bile components may be a plausible therapeutic approach to aging, improving liver fibrosis (41,42). In the MT staining results we obtained, a significant decrease in collagen density was detected in the treatment group given TUDCA. Our results are consistent with the studies mentioned above.

One process leading to age-related liver diseases is hepatic inflammation (16). Since inflammatory cytokines rise with hepatic aging, dysregulation of inflammasome function is a major factor in the uncontrolled immune response during inflammation (43). Hepatocytes are targeted by damaged cells, leading to the pathogenesis of age-related liver diseases (44). These age-related damages promote inflammation, and pro-inflammatory cytokines further potentiate inflammation (17). Activation of the NLRP3 inflammasome triggers different processes according to hepatocytes and other cell types (45). One of them is that the NLRP3 inflammasome contributes to the pathophysiology of age-related liver diseases and induces hepatocytes to form lipid droplets, which in turn promotes dyslipidemia (46). Previous studies have reported higher NLRP3 and ASC levels in liver samples due to aging (15,45,47,48). From the data obtained, it was concluded that the NLRP3 inflammasome plays a central role in determining the severity of liver inflammation. In the present study, SCD probiotics and TUDCA treatments were shown to regulate lipid metabolism and modulate NLRP3 inflammasome activation by reducing steatosis in the aged liver. Taken together, we report that the active role of the NLRP3 inflammasome in the relationship between the uncontrolled activation of the age-related immune response and the natural aging process, which may exacerbate liver inflammation and fibrosis, alleviates age-related chronic inflammatory hepatic pathology with anti-inflammatory therapies and may therefore be a target for age-related liver disease.

Low alanine aminotransferase (ALT) and aspartate aminotransferase (AST) levels in the low physiological range are surrogate markers for decreased liver metabolic function. These enzyme levels may suggest a relationship with other chronic diseases that occur during the aging process (49). In a previous study, it was reported that reductions in the aspartate-alanine aminotransferase ratio during the aging period can provide insight into its ability to predict nonalcoholic fatty liver disease and advanced fibrosis (50). In our study findings, we found that LDH levels decreased in the elderly control group and increased significantly in the groups given SCD Probiotics, TUDCA, and combined supplements. As muscle and bone mass decrease with age, this may cause a decrease in the LDH enzyme levels. Since SCD Probiotics and TUDCA supplements can accelerate metabolism, which is low in elderly individuals, increased metabolic activity may have increased the LDH enzyme level. The data we obtained are compatible with the data of Kaczor et al. (2006). Furthermore, lower LDH activity in older adults is a robust finding still present after accounting for the lower total protein content associated with aging.

## CONCLUSION

This study examined the combined impact of SCD Probiotics and TUDCA on the liver tissue of older male rats, offering insights into their potential therapeutic benefits. Using ATR-FTIR spectroscopy and machine learning, it explored changes in the liver at the molecular level, indicating both TUDCA and SCD Probiotics may alleviate age-related liver fibrosis and improve liver function. Notable changes in lipid, protein, and nucleic acid bands were observed, alongside significant improvements in liver fibrosis, cellular degradation, lymphatic infiltration, and bile duct growth. Furthermore, TUDCA and SCD Probiotics reduced liver microvesicular steatosis and collagen accumulation in older rats. Although all treatment groups saw increased serum AST and LDH levels, only the SCD Probiotics group exhibited a significant increase in albumin levels. SCD Probiotics exhibited a marked reduction in inflammasome-related gene expressions, and the lowest gene expression levels were observed in the group receiving both treatments This suggests different molecular mechanisms behind the positive effects of TUDCA and SCD Probiotics on liver health. The findings lay the groundwork for further research into the exact mechanisms through which these treatments benefit liver health. Understanding the effect of SCD probiotics and TUDCA supplements, NLRP3 inflammasome in ageing may pave the way to developing a targeted and adequate therapeutic strategy. Future studies should explore the long-term effects and the potential use of these treatments in clinical settings to prevent and treat age-related liver diseases.

### Author Contributions

HTT and TC contributed to all aspects of this study, from conceptualization to writing and editing. SK performed the histopathological and biochemical analysis, interpreted results, and contributed to writing and editing. BB, GS, HT were involved in the investigation, methodology, validation, and visualization. EA contributed to histopathological interpretation, visualization, and manuscript editing. RG conducted the investigation, methodology, and contributed to writing and editing. All authors have approved the final manuscript.

### Data Availability

All data generated during and/or analyzed during the current study are available from the corresponding author upon reasonable request.

### Ethical Approval

This study was approved by the Ethics Committee (approval number: 2022/03) of the Saki Yenilli Experimental Animal Production and Practice Laboratory.

### Funding

No financial support was received for this study.

