## Supplementary_Material_Methods for "Promoting Longevity in Aged Liver through NLRP3 Inflammasome Inhibition Using Tauroursodeoxycholic Acid (TUDCA) and SCD Probiotics"

### MATERIAL METHOD

#### Animal studies

The study used male Sprague-Dawley rats (24 months old) as the model organism. The animals were divided into four groups: the control group (n=7), the group that received probiotics (n=7), the group that was administered TUDCA (n=7), and the group that received TUDCA during SCD Probiotics supplementation (n=7), with each group receiving treatment for seven days. The rats were fed a standard rodent diet ad libitum throughout the study. TUDCA was administered intravenously at a dose of 300 mg/kg via the tail, while the probiotic supplement was given orally by gavage at a dose of 3 mL (1 x 108 CFU) per day. The study used a probiotic product containing 11 different probiotics marketed by SCD Probiotics company (Essential Probiotics XI - 500 ml H.S. Code: 2206.00.7000), including *Bacillus subtilis, Bifidobacterium bifidum, Bifidobacterium longum, Lactobacillus acidophilus, Lactobacillus bulgaricus, Lactobacillus casei, Lactobacillus fermentum, Lactobacillus plantarum, Lactococcus lactis, Saccharomyces cerevisiae,* and *Streptococcus thermophilus* species. After the treatment, the animals were euthanized one day later by ether treatment, and the liver tissues were extracted, immediately shocked on dry ice, and stored in a -80°C deep freezer until analysis. The animals were housed according to standard animal care protocols, and the study was approved by the Ethics Committee (approval number: 2022/03) of the Saki Yenilli Experimental Animal Production and Practice Laboratory.

#### Analysis of samples by Attenuated Total Reflectance Fourier Transform Infrared (ATR-FTIR) spectroscopy

Liver samples of all animals (2x24=48 in total) were compressed on the Zn/Se crystal of the ATR unit (PerkinElmer) without any pretreatment and examined with an ATR-FTIR spectrometer (PerkinElmer) at a resolution of 4 cm-1 and a scan number of 32. The spectra were obtained with the Spectrum One (PerkinElmer) software in the wavelength range of 4000-650 cm-1 (1).

**Prediction studies with different machine learning approaches based on big spectral data**

Linear Discriminant Analysis (LDA) and Support Vector Machine (SVM) were applied to differentiate the experimental groups using spectral data obtained from FTIR spectrometers. The spectral data were pre-processed using The Unscrambler® X 10.3 (CAMO Software AS, Norway) software with a baseline offset transformation in the 4000-650 $cm^{-1}$ region. The pre-processed spectra were first subjected to Principal Component Analysis (PCA), an unsupervised pattern processing technique. After standard deviation normalization (mean centering normalization) and validation using the leverage-correction method, the spectra were examined in lipid (3000-2700 cm-1), protein (1700-1500 cm-1), nucleic acid and polysaccharide (1200-650 cm-1), and full (4000-650 cm-1) regions by Singular Value Decomposition (SVD) algorithm (2).

LDA is a supervised classifier that linearly transforms n-dimensional feature samples into an m-dimensional space. In this study, PCA data were utilized as LDA model inputs with The Unscrambler® X 10.3 (CAMO Software AS, Norway) multivariate analysis (MVA) software. The category variable column was included in a data matrix, and all spectra of different sample categories were used to generate a training set. The quadratic method using the projections of the 7 PCA components was used for the prediction. Prior probabilities were calculated from the training set. The LDA results were presented as a discrimination plot, prediction matrix, and confusion matrix (3).

SVM is another general machine learning approach used in this study. The SVM classification method was accomplished using The Unscrambler® X 10.3 (CAMO Software

AS, Norway) multivariate analysis (MVA) software. All spectra were pre-processed, as explained earlier, and different sample categories were used to generate a training set. Classification (nu-SVC) was chosen as the SVM type using a linear method as the Kernel type. Nu value was set to 0.5, weights as all 1.00. The 14 segments of cross-validation were used in the calculation of training and cross-validation accuracies. Finally, the generated training dataset was applied to all sample datasets to obtain an SVM classification model (4).

**Quantification studies of FTIR spectral bands**

Spectral data analysis was performed using OPUS 5.5 (Bruker) software. The average spectra of each sample were baseline corrected using the Rubberband correction method with 64 baseline points before the band quantification analyses. In detailed band analyses, the bands with the highest absorbance values in different spectral regions of the spectra were selected, and the beginning and ending frequencies of the bands were determined with precision. The areas of bands specific to various biomolecules were analyzed by taking the integral areas of the determined frequency ranges with the OPUS 5.5 (Bruker) software. In addition, a virtual line was drawn vertically from the band baseline's midpoint to the band's peak, and the length of the line was measured with the help of a virtual ruler. Then, by marking the point where 0.75 times the line length coincides with the line, a horizontal line was drawn along the band from this point, and bandwidth values were obtained (5).

**Histopathological Examination**

**Hematoxylin & eosin (H&E) staining**

After the sacrification, liver samples were fixed in neutral buffered formalin (10%) for 48 h, followed histological routine tissue processing as in the previous study (6,8). Liver tissue was cut into 5 μm thick tissue sections from paraffin blocks using a rotary microtome (Leica

Biosystems, Germany). All slices were placed on glass slides, and stained with H&E, and viewed with an Olympus BX53 (Olympus, Tokyo, Japan) light microscope.

**Masson trichrome (MT) staining**

After deparafinization, other liver sections were used for MT staining according to the commercial kit (Cat No: 04-010802, Bio-Optica, Milan, Italy) following the instructions to assess liver fibrosis of collagen density for evaluating effects of TUDCA, SCD-Probiotics and combined treatment. All procedures were carried out at room temperature.

**Quantification of histological parameters**

H&E evaluations of lymphatic infiltration and micro vesicular steatosis (micro-lipid droplets) were performed using Image J (Fiji). The same method was applied for measuring MT-positive areas. This quantification method was adapted according to literature (7). Microscopic analysis was carried out at ×200 magnification, and the main features of hepatic fatty infiltration, and micro vesicular steatosis, were observed in the experimental groups. All stained areas below the same threshold were measured and signal intensities from five images in each section were plotted for all rats in the same group using an image processing program. 10-12 areas per animal section were randomly sampled for histopathological examination (6). All microscopic analyses were performed by two observers blinded to the study. Microphotographs were analyzed using light microscopy (Olympus BX53, Japan) with a camera attachment (Olympus DP27, Japan) and imaging systems (Olympus cellSens Entry, Japan) (8).

**Immunohistochemical analysis**

All procedure was processed according to the previously described protocol for paraffin sections to elucidate the possible effects of SCD probiotics and TUDCA on NLRP3 inflammasome (NLRP3, and ASC) on the aged liver tissue. Liver tissue sections were deparaffinized, rehydrated, and underwent endogenous peroxidase activity blocking solution for 10 min in hydrogen peroxide (3%) to inhibit reactions of non-specific antibody binding sites, then samples were washed with PBS. Furthermore, the sections subjected to 10 mM sodium citrate buffer to incubate heat-induced antigen retrieval (pH 6.0) for two or three consecutive runs in a microwave each for 4-5 min then left to cool down at room temperature. After the blocking step, the sections were incubated at 4°C overnight with anti-NLRP3 (Elabscience, E-AB-93112, dilution rate: 1:200), and anti-ASC (Elabscience, E-AB-30582, dilution rate: 1:200) polyclonal antibodies. The following day, the sections were subjected to a process involving biotinylated antibodies (TP-125-BN, Thermo Scientific) and streptavidin peroxidase (TS-125-HR). 3–3'Diaminobenzidine (DAB substrate kit, ab64238, Abcam) immunodetection was performed as a chromogen for 3–5 min. Sections were counterstained with Mayer's hematoxylin, dehydrated, and mounted. All images were taken of immune-stained areas using a camera (Olympus DP27, Japan) and imaging systems (Olympus cellSens Entry, Japan) under a light microscope (Olympus BX53, Japan). To quantify intensity, areas were randomly selected, and immunoprecipitation of stained cells was performed using Image J (Fiji). The intensities of NLRP3 and ASC expressions were performed in ten randomly selected fields for each group (3 different sections for each rat per group) (9).

**Biochemical analysis**

After collecting blood samples while all rats were under euthanasia, all samples were centrifugated for 15 minutes at 3500 rpm, and a refrigerated centrifuge was used to separate serum at 4 °C. Enzymatic assays were performed on a Hitachi C502 automated biochemical analyzer (Roche, Germany) with commercial kits (Roche Diagnostics) to measure serum levels of aspartate aminotransferase (AST), alanine aminotransferase (ALT), alkaline phosphatase (ALP), lactate dehydrogenase (LDH), and albumin, which were then expressed as IU/mL and g/L, respectively. All parameters were carried out in accordance with the protocols that were provided by the manufacturer (Roche Diagnostics).

### Statistics

Statistical evaluations and graph plots of the results were made using GraphPad Prism 9.01 (GraphPad, USA). The data were analyzed using One-way ANOVA and/or unpaired t-test, and the significance levels were stated as P 0.05 *, $P \leq 0.01$ **, $P \leq 0.001$ ***, and $P \leq 0.0001$ ****. Results are presented as mean ± SEM (standard error of the mean).
