## Supplementar_Figures for "Promoting Longevity in Aged Liver through NLRP3 Inflammasome Inhibition Using Tauroursodeoxycholic Acid (TUDCA) and SCD Probiotics"

### Supplementary Figures

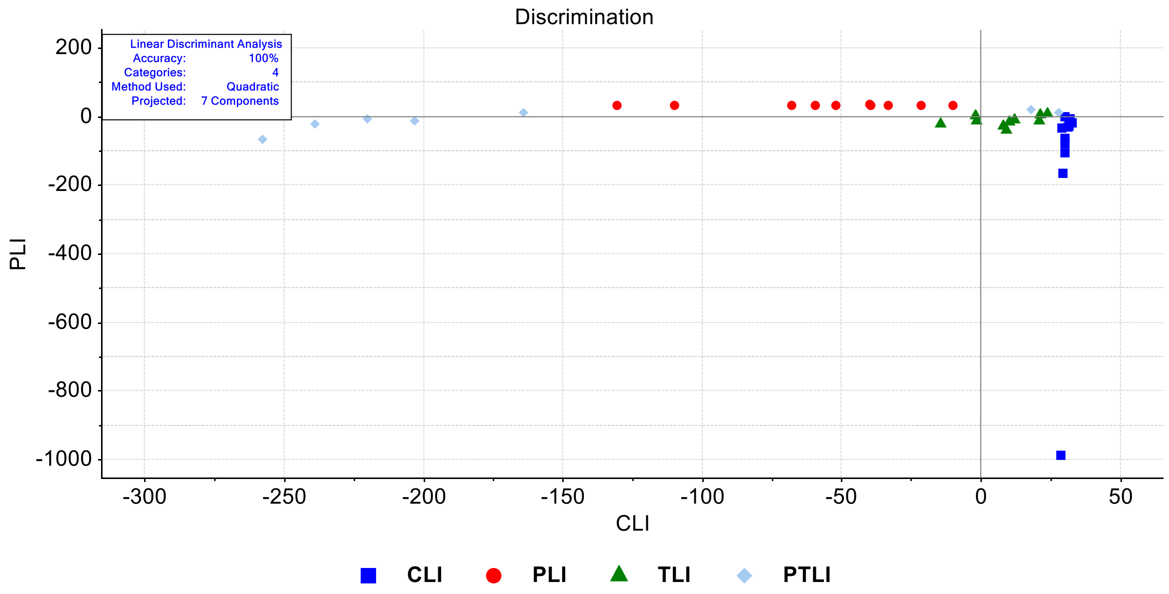


**Figure S1.** LDA discrimination plot for liver samples in the protein (1700-1500 $cm^{-1}$) spectral region. CLI (control), TLI (TUDCA)**,** PLI (SDC Probiotics), and the PTLI applications (in which the TUDCA and SCD Probiotics were applied together)

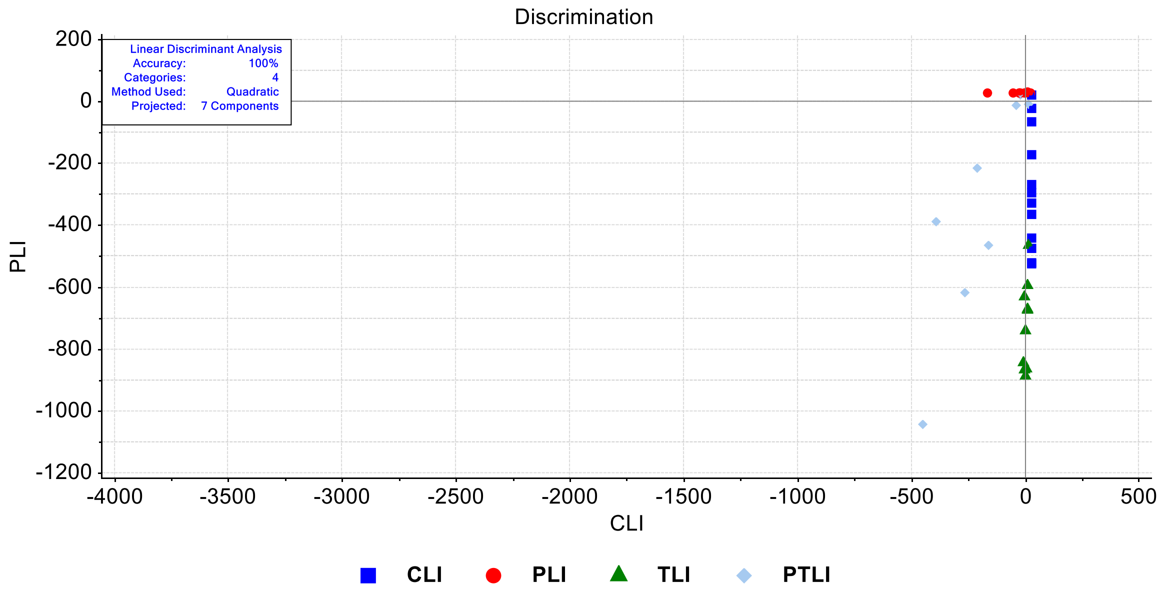


**Figure S2.** LDA discrimination plot for liver samples in the spectral region of the nucleic acid and polysaccharide (1200-650 $cm^{-1}$). CLI (control), TLI (TUDCA), PLI (SDC Probiotics), and the PTLI applications (in which the TUDCA and SCD Probiotics were applied together)

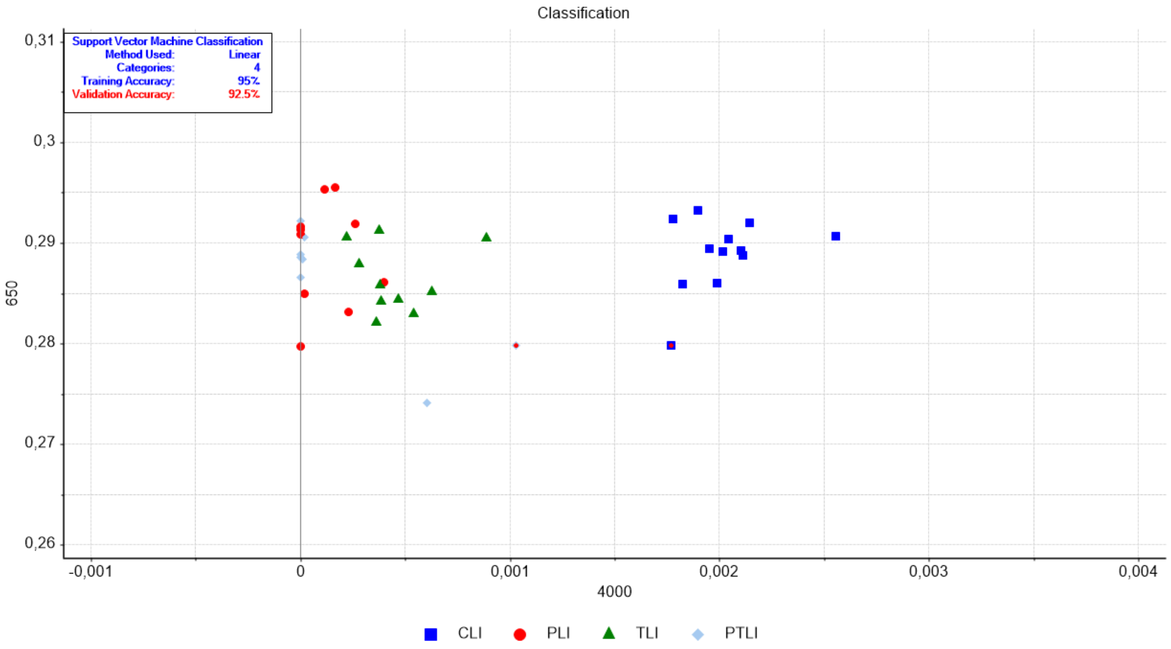


**Figure S3.** SVM classification plot for liver samples in the full (4000-650 $cm^{-1}$) spectral region. CIL (control), TIL (TUDCA)**,** PIL (SDC Probiotics), and the PTIL applications (in which the TUDCA and SCD Probiotics were applied together)

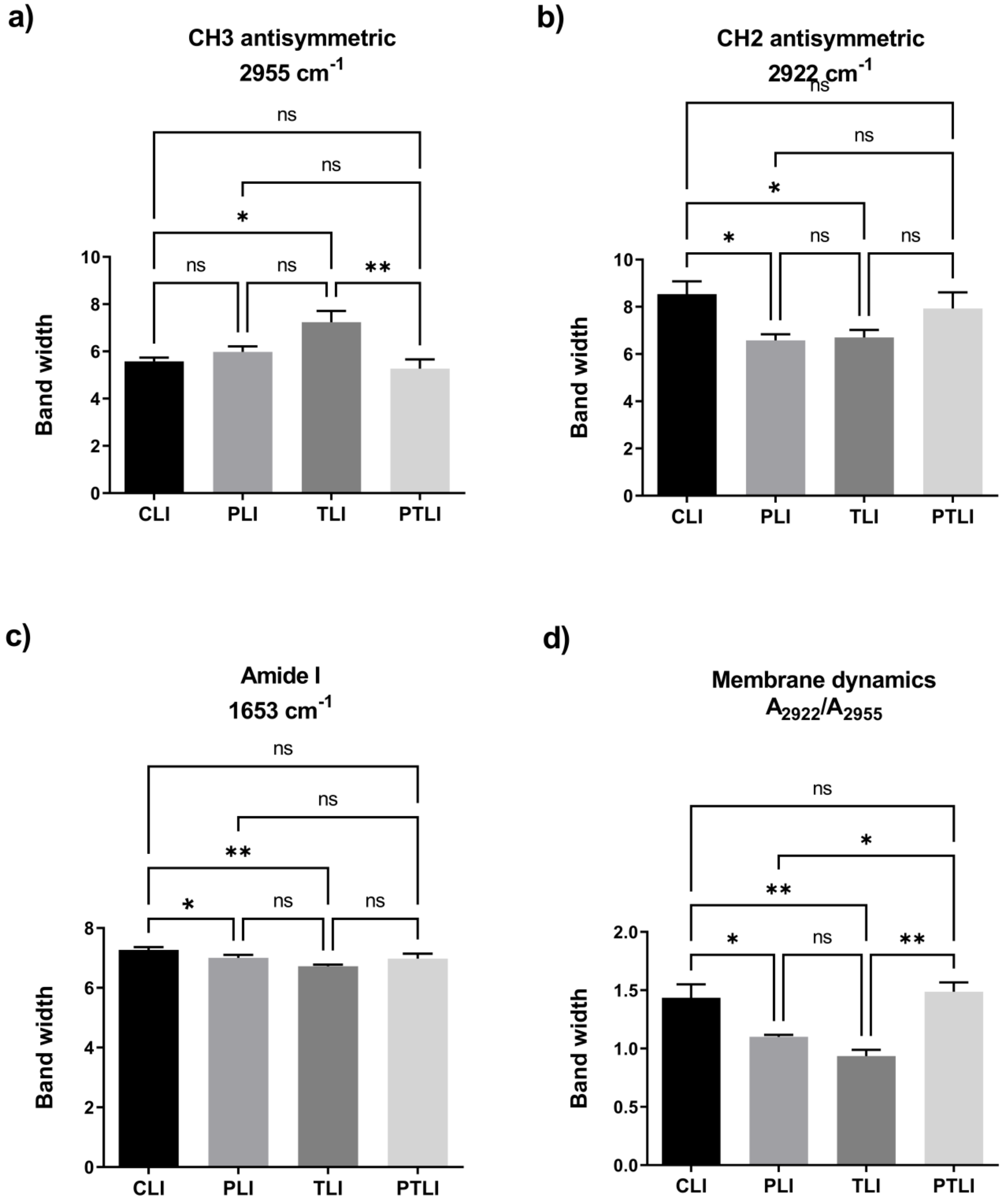


**Figure S4.** The quantitative changes in liver-associated spectrochemical parameters. The bandwidths for **a)** $CH_3$ antisymmetric (2955 $cm^{-1}$), **b)** $CH_2$ antisymmetric (2922 $cm^{-1}$), **c)** Amide I (1653 $cm^{-1}$), and **d)** membrane dynamics ($A_{2922}/A_{2955}$). CLI (control), TLI (TUDCA), PLI (SDC Probiotics), and the PTLI applications (in which the TUDCA and SCD Probiotics were applied together)
