## Supplementary_Tables for "Promoting Longevity in Aged Liver through NLRP3 Inflammasome Inhibition Using Tauroursodeoxycholic Acid (TUDCA) and SCD Probiotics"

**Table S1** LDA confusion matrix for liver samples in the full (4000-650 $cm^{-1}$) spectral region. CLI (control), TLI (TUDCA), PLI (SDC Probiotics), and the PTLI applications (in which the TUDCA and SCD Probiotics were applied together)

| Confusion matrix | Actual | CLI | PLI | TLI | PTLI |
|---|---|---|---|---|---|
| Predicted | | 1 | 2 | 3 | 4 |
| **CLI** | 1 | **12** | 0 | 0 | 0 |
| **PLI** | 2 | 0 | **10** | 0 | 0 |
| **TLI** | 3 | 0 | 0 | **10** | 0 |
| **PTLI** | 4 | 0 | 0 | 0 | **8** |

**Table S2** LDA prediction matrix for liver samples in the full (4000-650 $cm^{-1}$) spectral region. CLI (control), TLI (TUDCA), PLI (SDC Probiotics), and the PTLI applications (in which the TUDCA and SCD Probiotics were applied together)

| | CLI | PLI | TLI | PTLI | Predicted |
|---|---|---|---|---|---|
| | 1 | 2 | 3 | 4 | 5 |
| 1 | 18,62 | -277,92 | -435,14 | -203,15 | **CLI** |
| 2 | 19,70 | -54,27 | -557,18 | -875,02 | **CLI** |
| 3 | 20,93 | -20,48 | -488,99 | -1536,03 | **CLI** |
| 4 | 21,21 | -18,84 | -375,06 | -1641,06 | **CLI** |
| 5 | 20,75 | -164,16 | -297,49 | -1654,43 | **CLI** |
| 6 | 20,72 | -296,59 | -86,43 | -2488,77 | **CLI** |
| 7 | 21,18 | -49,52 | -690,34 | -366,65 | **CLI** |
| 8 | 20,42 | -64,44 | -542,97 | -1021,44 | **CLI** |
| 9 | 19,84 | -165,34 | -453,36 | -198,71 | **CLI** |
| 10 | 21,67 | -86,29 | -395,53 | -1039,68 | **CLI** |
| 11 | 18,71 | -361,10 | -313,45 | -289,42 | **CLI** |
| 12 | 20,84 | -189,89 | -228,08 | -1363,39 | **CLI** |
| 13 | -66,66 | 21,35 | -1408,25 | -3247,80 | **PLI** |
| 14 | -24,12 | 21,64 | -660,80 | -1667,07 | **PLI** |
| 15 | -3,70 | 21,20 | -541,61 | -404,12 | **PLI** |

| | | | | | |
|---|---|---|---|---|---|
| 16 | -15,73 | 21,99 | -416,60 | -345,50 | **PLI** |
| 17 | -27,96 | 21,68 | -520,57 | -1141,84 | **PLI** |
| 18 | -68,29 | 20,75 | -502,50 | -1228,19 | **PLI** |
| 19 | -46,40 | 23,57 | -401,53 | -543,61 | **PLI** |
| 20 | -117,94 | 21,18 | -193,66 | -428,24 | **PLI** |
| 21 | -56,31 | 20,77 | -275,92 | -1170,24 | **PLI** |
| 22 | -116,42 | 20,96 | -178,19 | -1641,27 | **PLI** |
| 23 | -82,76 | -494,45 | 25,17 | -2754,42 | **TLI** |
| 24 | -138,40 | -865,97 | 24,81 | -3150,50 | **TLI** |
| 25 | -55,55 | -426,82 | 24,57 | -3619,71 | **TLI** |
| 26 | -147,42 | -779,84 | 24,33 | -4112,47 | **TLI** |
| 27 | -90,08 | -493,96 | 26,04 | -4270,52 | **TLI** |
| 28 | -106,08 | -474,39 | 26,34 | -4170,08 | **TLI** |
| 29 | -72,64 | -456,06 | 25,89 | -3254,97 | **TLI** |
| 30 | -80,95 | -542,63 | 24,28 | -2681,58 | **TLI** |
| 31 | -160,88 | -464,42 | 24,35 | -1880,94 | **TLI** |
| 32 | -154,19 | -647,74 | 24,69 | -3103,25 | **TLI** |
| 33 | 2,41 | 0,12 | -470,33 | 21,77 | **PTLI** |
| 34 | -11,57 | -19,14 | -432,47 | 21,77 | **PTLI** |
| 35 | -98,57 | -101,98 | -112,00 | 21,77 | **PTLI** |
| 36 | -213,60 | -358,42 | -231,01 | 21,77 | **PTLI** |
| 37 | -309,16 | -897,82 | -354,61 | 21,77 | **PTLI** |
| 38 | -376,98 | -764,30 | -256,69 | 21,77 | **PTLI** |
| 39 | -398,05 | -785,97 | -351,88 | 21,77 | **PTLI** |
| 40 | -381,32 | -1044,74 | -429,09 | 21,77 | **PTLI** |

**Table S3** LDA confusion matrix for liver samples in the lipid (3000-2700 $cm^{-1}$) spectral region. CLI (control), TLI (TUDCA)**,** PLI (SDC Probiotics), and the PTLI applications (in which the TUDCA and SCD Probiotics were applied together)

| Confusion matrix | Actual | CLI | PLI | TLI | PTLI |
|---|---|---|---|---|---|
| Predicted | | 1 | 2 | 3 | 4 |
| **CLI** | 1 | **12** | 0 | 0 | 0 |
| **PLI** | 2 | 0 | **10** | 0 | 0 |
| **TLI** | 3 | 0 | 0 | **10** | 0 |
| **PTLI** | 4 | 0 | 0 | 0 | **8** |

**Table S4** LDA prediction matrix for liver samples in the lipid (3000-2700 $cm^{-1}$) spectral region. CLI (control), TLI (TUDCA)**,** PLI (SDC Probiotics), and the PTLI applications (in which the TUDCA and SCD Probiotics were applied together)

| | CLI | PLI | TLI | PTLI | Predicted |
|---|---|---|---|---|---|
| | 1 | 2 | 3 | 4 | 5 |
| 1 | 40,19 | -98,56 | -2145,10 | -19,63 | **CLI** |
| 2 | 41,27 | -17,84 | -2269,72 | -19,74 | **CLI** |
| 3 | 42,85 | -1,81 | -1642,60 | 34,76 | **CLI** |
| 4 | 42,99 | -3,59 | -1324,66 | 34,32 | **CLI** |
| 5 | 40,73 | 1,32 | -2020,81 | -1,70 | **CLI** |
| 6 | 41,11 | -8,40 | -1224,53 | -27,40 | **CLI** |
| 7 | 42,63 | -25,79 | -2430,00 | -13,47 | **CLI** |
| 8 | 44,00 | -42,20 | -2102,07 | -18,59 | **CLI** |
| 9 | 42,45 | -28,42 | -2325,03 | -84,42 | **CLI** |
| 10 | 43,96 | -36,06 | -1845,10 | -52,42 | **CLI** |
| 11 | 40,17 | -112,78 | -92,95 | -134,96 | **CLI** |
| 12 | 41,36 | -45,53 | -1170,76 | -73,97 | **CLI** |
| 13 | -239,84 | 40,39 | -5604,29 | -230,88 | **PLI** |
| 14 | -82,74 | 41,47 | -3047,65 | 25,06 | **PLI** |
| 15 | -284,01 | 41,45 | -5214,07 | -87,98 | **PLI** |
| 16 | -339,93 | 40,48 | -5324,10 | -28,15 | **PLI** |
| 17 | -499,56 | 40,43 | -6190,81 | -58,18 | **PLI** |
| 18 | -386,19 | 43,04 | -6265,92 | -107,10 | **PLI** |

| | | | | | |
|---|---|---|---|---|---|
| 19 | -423,80 | 42,78 | -5868,61 | -117,06 | **PLI** |
| 20 | -589,29 | 40,78 | -4600,51 | -89,25 | **PLI** |
| 21 | -557,67 | 40,71 | -6360,44 | -82,16 | **PLI** |
| 22 | -751,76 | 40,55 | -5772,57 | -96,85 | **PLI** |
| 23 | -81,63 | -37,32 | 48,33 | -84,61 | **TLI** |
| 24 | -111,89 | -81,04 | 47,29 | -29,35 | **TLI** |
| 25 | -21,37 | -22,32 | 47,27 | -80,91 | **TLI** |
| 26 | -118,62 | -53,28 | 47,25 | -31,24 | **TLI** |
| 27 | -87,34 | -40,41 | 47,93 | -70,85 | **TLI** |
| 28 | -75,55 | -39,34 | 49,10 | -85,88 | **TLI** |
| 29 | -68,06 | -25,79 | 47,89 | -110,37 | **TLI** |
| 30 | -85,07 | -33,89 | 47,39 | -91,32 | **TLI** |
| 31 | -81,90 | -63,83 | 48,43 | -38,12 | **TLI** |
| 32 | -91,40 | -66,44 | 49,73 | -36,47 | **TLI** |
| 33 | 29,11 | 11,36 | -1753,90 | 41,56 | **PTLI** |
| 34 | 27,08 | 28,20 | -2117,20 | 41,56 | **PTLI** |
| 35 | -83,16 | -53,84 | -1019,73 | 41,56 | **PTLI** |
| 36 | -70,20 | -508,55 | -1071,49 | 41,56 | **PTLI** |
| 37 | -51,68 | -466,85 | -793,60 | 41,56 | **PTLI** |
| 38 | -143,75 | -443,40 | -291,49 | 41,56 | **PTLI** |
| 39 | -96,30 | -519,10 | -1349,88 | 41,56 | **PTLI** |
| 40 | -163,59 | -939,20 | -902,47 | 41,56 | **PTLI** |

**Table S5** LDA confusion matrix for liver samples in the spectral region of the protein (1700-1500 cm-1). CLI (control), TLI (TUDCA), PLI (SDC Probiotics), and the PTLI applications (in which the TUDCA and SCD Probiotics were applied together)

| Confusion matrix | Actual | CLI | PLI | TLI | PTLI |
|---|---|---|---|---|---|
| Predicted | | 1 | 2 | 3 | 4 |
| CLI | 1 | **12** | 0 | 0 | 0 |
| PLI | 2 | 0 | **10** | 0 | 0 |
| TLI | 3 | 0 | 0 | **10** | 0 |
| PTLI | 4 | 0 | 0 | 0 | **8** |

**Table S6** LDA prediction matrix for liver samples in the spectral region of the protein (1700-1500 cm-1). CLI (control), TLI (TUDCA), PLI (SDC Probiotics), and the PTLI applications (in which the TUDCA and SCD Probiotics were applied together)

| | CLI | PLI | TLI | PTLI | Predicted |
|---|---|---|---|---|---|
| | 1 | 2 | 3 | 4 | 5 |
| 1 | 28,91 | -33,79 | -170,50 | -3313,31 | CLI |
| 2 | 30,26 | -0,57 | -3,74 | -495,88 | CLI |
| 3 | 31,44 | -31,23 | -7,08 | -14,69 | CLI |
| 4 | 31,74 | -27,27 | 0,82 | -43,21 | CLI |
| 5 | 30,06 | -78,52 | -173,59 | -301,13 | CLI |
| 6 | 29,55 | -164,89 | -342,63 | -2327,41 | CLI |
| 7 | 31,52 | -15,18 | 4,62 | -86,85 | CLI |
| 8 | 32,04 | -8,71 | 13,22 | -674,82 | CLI |
| 9 | 30,32 | -65,43 | -145,46 | -10,11 | CLI |
| 10 | 32,92 | -18,98 | 6,74 | -612,88 | CLI |
| 11 | 28,84 | -986,42 | -3287,17 | -2205,90 | CLI |
| 12 | 30,26 | -106,71 | -137,21 | -323,40 | CLI |
| 13 | -51,98 | 32,14 | -29,42 | -3232,26 | PLI |
| 14 | -39,58 | 32,90 | -58,08 | 14,46 | PLI |
| 15 | -21,59 | 32,93 | -0,95 | -832,44 | PLI |
| 16 | -33,38 | 31,93 | -2,45 | -1549,68 | PLI |
| 17 | -67,83 | 31,89 | -71,25 | -203,22 | PLI |
| 18 | -39,79 | 33,52 | -45,22 | -283,54 | PLI |

| | | | | | |
|---|---|---|---|---|---|
| 19 | -109,79 | 31,85 | -35,22 | -429,53 | **PLI** |
| 20 | -130,61 | 31,48 | -34,82 | -1350,44 | **PLI** |
| 21 | -9,92 | 33,08 | -39,72 | -574,33 | **PLI** |
| 22 | -59,31 | 31,74 | -237,63 | -1498,81 | **PLI** |
| 23 | 10,45 | -15,59 | 35,28 | -1735,36 | **TLI** |
| 24 | -14,24 | -22,97 | 33,84 | -5741,52 | **TLI** |
| 25 | 12,15 | -9,40 | 33,90 | -5936,62 | **TLI** |
| 26 | -1,47 | -13,57 | 33,56 | -4966,95 | **TLI** |
| 27 | 8,16 | -29,33 | 35,79 | -4877,67 | **TLI** |
| 28 | 9,18 | -39,72 | 35,07 | -4197,54 | **TLI** |
| 29 | 21,17 | 4,95 | 34,89 | -588,91 | **TLI** |
| 30 | 24,05 | 6,74 | 34,16 | -1958,34 | **TLI** |
| 31 | -1,83 | 2,41 | 33,82 | -773,42 | **TLI** |
| 32 | 20,79 | -14,64 | 34,27 | -502,97 | **TLI** |
| 33 | 28,15 | 10,92 | 23,34 | 33,26 | **PTLI** |
| 34 | 18,01 | 18,74 | -5,69 | 33,26 | **PTLI** |
| 35 | -250,87 | 1,85 | -279,93 | 33,26 | **PTLI** |
| 36 | -238,98 | -21,81 | -475,97 | 33,26 | **PTLI** |
| 37 | -220,22 | -7,53 | -631,89 | 33,26 | **PTLI** |
| 38 | -163,97 | 11,91 | -256,11 | 33,26 | **PTLI** |
| 39 | -203,28 | -12,84 | -350,70 | 33,26 | **PTLI** |
| 40 | -257,52 | -68,00 | -501,57 | 33,26 | **PTLI** |

**Table S7** LDA confusion matrix for liver samples in the spectral region of nucleic acids and polysaccharides (1200-650 $cm^{-1}$). CLI (control), TLI (TUDCA), PLI (SDC Probiotics), and the PTLI applications (in which the TUDCA and SCD Probiotics were applied together)

| Confusion matrix | Actual | CLI | PLI | TLI | PTLI |
|---|---|---|---|---|---|
| Predicted | | 1 | 2 | 3 | 4 |
| **CLI** | 1 | **12** | 0 | 0 | 0 |
| **PLI** | 2 | 0 | **10** | 0 | 0 |
| **TLI** | 3 | 0 | 0 | **10** | 0 |
| **PTLI** | 4 | 0 | 0 | 0 | **8** |

**Table S8** LDA prediction matrix for liver samples in the spectral region of nucleic acids and polysaccharides (1200-650 $cm^{-1}$). CLI (control), TLI (TUDCA), PLI (SDC Probiotics), and the PTLI applications (in which the TUDCA and SCD Probiotics were applied together)

| | CLI | PLI | TLI | PTLI | Predicted |
|---|---|---|---|---|---|
| | 1 | 2 | 3 | 4 | 5 |
| 1 | 27,22 | -172,21 | -517,36 | -327,13 | **CLI** |
| 2 | 28,02 | -65,02 | -125,13 | -331,32 | **CLI** |
| 3 | 29,03 | 22,10 | -1423,02 | -401,04 | **CLI** |
| 4 | 30,06 | -23,65 | -1007,16 | -431,55 | **CLI** |
| 5 | 29,37 | -295,95 | -1807,11 | -1359,35 | **CLI** |
| 6 | 29,95 | -473,64 | -2513,75 | -1659,08 | **CLI** |
| 7 | 31,22 | -269,08 | -374,00 | -603,06 | **CLI** |
| 8 | 28,21 | -363,65 | -306,72 | -635,94 | **CLI** |
| 9 | 30,28 | -442,35 | -770,07 | -1222,21 | **CLI** |
| 10 | 29,24 | -521,07 | -171,67 | -826,76 | **CLI** |
| 11 | 27,42 | -327,16 | -5610,38 | -1433,37 | **CLI** |
| 12 | 28,36 | -525,38 | -1189,52 | -1494,24 | **CLI** |
| 13 | -3834,20 | 27,55 | -26517,20 | -3140,44 | **PLI** |
| 14 | -24,48 | 28,91 | -88,59 | -40,16 | **PLI** |
| 15 | 26,86 | 28,23 | -592,08 | -64,96 | **PLI** |
| 16 | 2,32 | 29,28 | -894,16 | 23,36 | **PLI** |
| 17 | -24,40 | 28,77 | -1300,58 | -2,00 | **PLI** |
| 18 | -4,57 | 27,59 | -497,54 | -32,12 | **PLI** |

| | | | | | |
|---|---|---|---|---|---|
| 19 | 10,67 | 29,79 | -1283,77 | -141,54 | **PLI** |
| 20 | -19,65 | 28,15 | -1807,43 | -77,03 | **PLI** |
| 21 | -51,47 | 27,83 | -1288,23 | 2,00 | **PLI** |
| 22 | -164,38 | 28,18 | -2569,68 | 8,14 | **PLI** |
| 23 | 15,69 | -465,02 | 32,54 | -279,95 | **TLI** |
| 24 | 10,37 | -674,50 | 32,63 | -1137,84 | **TLI** |
| 25 | -1,09 | -632,02 | 32,43 | -435,64 | **TLI** |
| 26 | 6,18 | -865,38 | 32,50 | -901,34 | **TLI** |
| 27 | 12,11 | -672,72 | 33,76 | -464,88 | **TLI** |
| 28 | 12,22 | -595,15 | 34,11 | -420,45 | **TLI** |
| 29 | -6,54 | -846,32 | 34,12 | -337,02 | **TLI** |
| 30 | 2,74 | -887,55 | 33,76 | -455,79 | **TLI** |
| 31 | 2,53 | -740,19 | 33,38 | -701,25 | **TLI** |
| 32 | -2,97 | -869,03 | 33,02 | -750,97 | **TLI** |
| 33 | 17,20 | -8,79 | -409,21 | 29,33 | **PTLI** |
| 34 | -18,82 | 19,60 | -625,48 | 29,33 | **PTLI** |
| 35 | -37,64 | -12,31 | -941,89 | 29,33 | **PTLI** |
| 36 | -392,35 | -387,55 | -1224,18 | 29,33 | **PTLI** |
| 37 | -211,89 | -216,48 | -808,92 | 29,33 | **PTLI** |
| 38 | -161,68 | -463,47 | -378,79 | 29,33 | **PTLI** |
| 39 | -263,65 | -617,78 | -315,36 | 29,33 | **PTLI** |
| 40 | -448,33 | -1045,97 | -528,06 | 29,33 | **PTLI** |
